# The Neural Architecture of Dream Recall Frequency: Insights from Interindividual Variations in Brain Structure and Function

**DOI:** 10.1101/2025.05.20.655114

**Authors:** Mariana Pereira, Paola Galdi, Ting Mei, Renate Rutiku, Irene Klærke Mikkelsen, Alberto Llera, René Scheeringa, Christian Beckmann, Florian Krause, Martin Dresler, Kristian Sandberg

**Affiliations:** Donders Institute of Cognition and Behaviour, Radboud University Medical Center, Nijmegen, The Netherlands; School of Informatics, University of Edinburgh, Edinburgh, UK; Consciousness Lab, Institute of Psychology, Jagiellonian University, Krakow, Poland; Center of Functionally Integrative Neuroscience, Aarhus University, Aarhus, Denmark; Lis Data Solutions, Santander, Spain; Neurobiology Research Unit, Copenhagen University Hospital Rigshospitalet, Copenhagen, Denmark

**Author notes:** Corresponding author:* Mariana Pereira. Shared senior authorship (Florian Krause, Martin Dresler, Kristian Sandberg).

**Keywords:** dream recall frequency, dream traits, dreaming, neuroimaging

## Abstract

Dreaming represents a complex and universal aspect of human sleep, yet it remains an intriguing phenomenon, with the neural mechanisms underlying dream experiences and their frequency not fully understood. This study employs a multimodal neuroimaging approach, integrating quantitative multi-parameter mapping, diffusion tensor imaging, and resting-state functional MRI, to investigate the neural correlates of dream recall frequency (DRF) in a large cohort of 258 healthy individuals. By employing Linked Independent Component Analysis (LICA), we were able to discern distinctive patterns of brain structure and function that correlated with variations in DRF. Our findings elucidate a complex relationship between dream recall and brain microstructure integrity, particularly in white matter regions of the orbitofrontal cortex, parahippocampal gyrus, superior parietal lobule, and occipital cortex. Higher DRF was related to increased white matter microstructure integrity in these regions and decreased gray matter volume in occipital and temporal areas. In terms of functional measures, higher DRF was associated with reduced connectivity across a range of resting-state networks, including the default mode, visual, and dorsal attention networks. This was particularly evident in the right precuneus and posterior cingulate cortex. These results suggest that enhanced dream recall may be related to the organization of higher-order visual and cognitive processing areas, supporting a top-down model of dreaming. This study contributes to a more comprehensive understanding of the neural substrates underlying individual differences in dream recall, offering a foundation for future investigations into the neurobiology and causal relationships of dreaming.

## Introduction

Humans spend approximately one-third of their lives asleep, with a considerable proportion of this time dedicated to dreaming (Simor et al., 2022). Although it is a common experience for many, dreaming encompasses a number of complex processes that remain largely unknown to scientists. Firstly, regardless of specific brain physiology and connectivity during different stages of sleep, content-specific regions in posterior cortical areas are activated, thereby resulting in a dream experience (Siclari et al., 2017; Cataldi et al., 2024). Nevertheless, the occurrence of a dream does not necessarily guarantee its recall upon awakening. For a dream to be remembered, it must undergo successful encoding, whereby the experience is transformed into a lasting memory trace, and then retrieved upon waking (Nemeth, 2023). Numerous hypotheses regarding the potential functions of dreams exist (Revonsuo, 2000). Yet, testing them empirically is challenging, particularly due to the lack of a physiological marker for dreaming and the reliance on subjective dream reports as the primary method for accessing oneiric experiences. Beyond exploring the neural correlates of dreaming while they are happening, investigating dream traits such as dream recall frequency (DRF) offers insight into the intricate processes that contribute to the phenomenon of dreams (Schredl & Montasser, 1996). Although retrospective dream recall has limitations, including the potential biases of self-report scales and the fragility of memory that can lead to false recollections (Beaulieu-Prévost & Zadra, 2015), it remains the most efficient and cost-effective method for studying trait dream recall. Furthermore, in contrast to the practice of clustering participants into predefined low- and high-dream recall groups, an analysis of the full distribution of recall frequency can elucidate the anatomical and functional variations in the brain that underlie normal inter-individual differences in DRF rather than just the extremes of the spectrum. Here, we employ a data-driven approach integrating multiple neuroimaging modalities in light of existing knowledge on dream generation and recall mechanisms.

Lesion studies and electrophysiological research have identified specific brain regions and neural oscillations associated with dream experiences, yet the precise roles these brain areas play in the generation, encoding, and retrieval of dreams remain unclear. The global cessation of dreaming has been associated with lesions in or near the temporal-occipital-parietal junction, posterior cortical regions, and ventromedial prefrontal areas, either unilaterally or bilaterally (Solms, 2000). Conversely, lesions in the prefrontal and anterior cingulate cortices have been linked to an increase in the frequency of dreams, as well as an increase in dream vividness and dream reality confusion (Solms, 2000; Vallat et al., 2018). From an electrophysiological perspective, dream experiences during both non-rapid eye movement (NREM) and REM sleep exhibit common features. Local high-frequency (20–50 Hz) activity over the “posterior hot zone” correlates with dream content, while increased high-frequency activity over medial and lateral frontal areas is associated with memory formation and storage (Siclari et al., 2017). Among the various theories regarding the production and functions of dreams, certain aspects of this complex process may influence the extent to which a dream experience is successfully recalled. The occurrence and intensity of the dream, local brain activations, and post-awakening interferences may all be factors in determining whether a dream is recalled or not (for a detailed review of these factors, see (Nemeth, 2023)).

Considerable inter-individual variation in DRF is influenced by a range of behavior and cognitive factors that seem to be intricately linked to structural and functional brain differences. In the general healthy population, this variability has been associated with a number of individual factors, including age, gender, personality traits, sleep habits, visual imagery, and creativity (Schredl & Montasser, 1996). Furthermore, studies examining the relationship between DRF and neuroimaging have revealed a complex interplay of structural and functional brain differences contributing to individual variations. For instance, neuroanatomical measures of deep gray matter structures such as the amygdala and hippocampus are not associated with DRF per se. However, they relate to qualitative aspects of dreams, including length, emotional load, bizarreness, and vividness (De Gennaro et al., 2011). Individuals with high DRF demonstrate higher regional cerebral blood flow (rCBF) in the temporoparietal junction during REM sleep, NREM stage 3, and wakefulness, as well as in the medial prefrontal cortex during REM sleep and wakefulness. No significant differences were reported in the medial prefrontal cortex during NREM stages 2 and 3, and no behavioral or cognitive differences were identified between groups (Eichenlaub et al., 2014). A negative correlation was observed between DRF and cortical volume in the medial fusiform and parahippocampal gyrus in the right hemisphere but not in the left. White matter integrity in fibers connected to these regions, particularly in the fusiform gyrus and inferior longitudinal fasciculus, negatively correlates with DRF (Zhou et al., 2019). Another MRI study found no significant differences in grey matter density between high and low recallers. However, an increase in white matter density in the medial prefrontal cortex of high recallers was observed, suggesting a potential role in dream production (Vallat et al., 2018). In terms of functional measures, DRF is negatively correlated with connectivity in a number of networks, including the visual, thalamic, basal ganglia, and auditory networks. Of particular note are the lateral visual network during the night and the posterior cingulate cortex in the morning (Zou et al., 2018). These findings highlight the complex and multifaceted relationship between DRF and a range of neuroimaging measures. They also suggest that both structural and functional brain differences contribute to individual differences in dream recall. However, the shared relationship across different neuroimaging modalities remains to be explored.

Linked independent component analysis (LICA) is a refined multimodal data fusion technique that simultaneously analyses multiple neuroimaging modalities, such as structural Magnetic Resonance Imaging (MRI), functional MRI, and diffusion tensor imaging (DTI), with the objective of identifying independent patterns of shared variance across these modalities (Groves et al., 2011; Llera et al., 2019). This method integrates input data at an early stage of the analysis pipeline rather than combining unimodal results post hoc, resulting in a more holistic understanding of brain-behavior relationships. LICA has been effectively utilized to elucidate the underlying neurobiology of several neurodevelopmental disorders, including autism spectrum disorder (Mei et al., 2023; Van Oort et al., 2023), obsessive-compulsive disorder (Xu et al., 2024), and attention deficit hyperactivity disorder (Itahashi et al., 2015), as well as demographic and behavioral characteristics (Llera et al., 2019; Kohn et al., 2021). The main advantage of LICA is its ability to enhance robustness to noise and its sensitivity to detect subtle effects in high-dimensional data that may be overlooked by univariate approaches. This is achieved by leveraging the complementary aspects of each imaging modality and efficiently modelling the shared variance. Moreover, LICA enables the investigation of inter-individual differences in brain measures and their relationships to behavioral and clinical phenotypes, which can offer insights into conventional diagnostic procedures. Additionally, it is emerging as a powerful tool for advancing our understanding of the complex interactions between brain structure, function, and behavior in both specific and transdiagnostic contexts.

This study leverages the power of this novel method to investigate the relationship between brain structural and functional characteristics with individual variations in DRF in a large dataset. We employed quantitative multi-parameter mapping and DTI to examine gray and white matter volume and morphology, respectively, and resting-state functional MRI to assess brain connectivity patterns associated with DRF. This comprehensive approach enabled the identification of potential anatomical and functional correlates of DRF, thereby providing a more nuanced understanding of the neural mechanisms underlying dream generation. By applying LICA to a large cohort of over 250 healthy individuals, we aimed to investigate the integrated structural and functional brain patterns that differentiate the full frequency spectrum of dream recall. This approach contributes to a broader understanding of how individual neurobiological variations influence DRF and the generation of dream experiences.

## Methods

The data utilized in this study is part of a large, multi-site study under the EU COST Action CA18106 (The Neural Architecture of Consciousness). The dataset encompasses MRI and behavioral data collected from healthy participants. The local ethics committee, *De Videnskabsetiske Komitéer for Region Midtjylland*, Denmark, approved the research protocol. The participants were recruited through the Center of Functionally Integrative Neuroscience (Aarhus University) participant database and local advertisement. Some data from the overall project has been published in other articles with different aims, and parts of the methods descriptions have been adapted from these articles as well as manuscripts in preparation. Specifically, dream recall data has previously been used in an article focusing purely on behavioral analyses (Tzioridou et al., 2022), and as a control variable in a manuscript investigating nightmare frequency in the context of emotional regulation (Pereira et al., 2024).

### Participants

A total of 306 participants consented to participate in the study and were compensated financially for their time and contributions. Of the total number of participants, 269 had MRI data available, of which eleven participants were excluded: five due to incomplete questionnaires, three due to incomplete functional MRI data, and three due to poor structural MRI quality and excessive movement artifacts. Hence, the final sample consisted of 258 participants (152 female, with a mean age of 24.89 ranging from 18 to 48 years).

### Behavioral materials and procedure

All participants completed an online questionnaire session from home with a total duration of around 70 minutes, including a seven-point rating scale assessing their DRF (Schredl & Erlacher, 2004), and general health. Typically within a few weeks of the scans, in an optional session, they completed the Wechsler Adult Intelligence Scale, Fourth Edition (WAIS-IV) (Lichtenberger & Kaufman, 2012). The participants were instructed to ensure the questionnaires were completed in an undisturbed environment. The DRF scale was recoded into units of mornings per week (Stumbrys et al., 2015). Although the evidence for a direct association between DRF and IQ is inconclusive, there is a body of literature indicating a link between IQ and REM sleep density (Busby & Pivik, 1983). Therefore, we sought to adjust for this potential confounding variable in our analysis. Because thirty-one participants did not complete the WAIS-IV questionnaire, missing data were handled using mean imputation, an approach that is appropriate for datasets where missing values are considered to be missing completely at random (Rubin, 2004). While mean imputation is a simple method, it can reduce variability in the data and avoid decreasing the sample size.

### MRI data acquisition

The imaging procedures were performed using a Siemens Magnetom Prisma-fit 3T MRI scanner. Two resting-state fMRI runs (12 and 6 minutes) were recorded alongside quantitative multi-parameter mapping (MPM; (Weiskopf et al., 2013)) and diffusion-weighted imaging in an approximately one-hour scanning session. For each participant, 1500 functional volumes were acquired using an echo planar T2*-weighted sequence sensitive to blood-oxygen-level-dependent (BOLD) contrast with a multiband acceleration factor of 6 (TR = 700 ms; TE = 33 ms; flip-angle = 53°, field of view = 200 × 200 mm, number of slices = 60; slice thickness = 2.5 mm [no gap]; in-plane resolution = 2.5 × 2.5 mm).

The MPM protocol was implemented based on the Siemens vendor sequence. Three-dimensional (3D) data acquisition consisted of three multi-echo spoiled gradient echo scans (i.e., fast low angle shot [FLASH] sequences with magnetization transfer saturation (MT), T1, and effective proton density (PD) contrast weighting). Additional reference radio-frequency (RF) scans were acquired. The acquisition protocol had the following parameters: TR = 18 ms (PDw/T1w) and 37 ms (MTw); TE = 2.46/4.92/7.38/9.84/12.30/14.76 ms (PDw/T1w/MTw); flip-angle = 6° (MTw), 4° (PDw), and 25° (T1w); voxel size = 1 mm^3^; field of view = 224 x 256 x 176 mm; phase encoding direction = AP; GRAPPA = 2; acquisition times = 3:50 (T1w/PDw) and 7:52 (MTw).

Diffusion-weighted imaging (dMRI) data were acquired using a High-angular resolution diffusion imaging (HARDI) protocol conducted within the same session, lasting approximately 10 minutes. The HARDI sequence encompassed multiple diffusion directions: 75 at b = 2500 s/mm^2^, 60 at b = 1500 s/mm^2^, 21 at b = 1200 s/mm^2^, 30 at b = 1000 s/mm^2^, 15 at b = 700 s/mm^2^, and 10 at b = 5 s/mm^2^. These varying b-shells were acquired in a single series with the following parameters: flip angle = 90°; TR = 2850ms; TE = 7 ms; voxel size = 2 mm^3^; matrix size of 100 x 100, and 84 slices; phase-encoding direction = AP with an additional acquisition in the opposite phase-encoding direction (PA) at b = 0, 700, 1000, 1200, 1500, 2500 s/mm^2^ for EPI distortion correction.

### Structural MRI data pre-processing and gray-matter volume estimation

Synthetic T1w images were generated using the longitudinal relaxation rate (R1) and effective proton density (PD) high-resolution maps (acquired during the MPM sequence protocol). First, both maps were thresholded to achieve the required FreeSurfer units. The R1 map was transformed into a T1 map by inverting its values, then thresholded at zero, and multiplied by one thousand to convert to milliseconds. The PD map was thresholded by zero and multiplied by one hundred. All manipulations were performed using *FSL maths* commands. Subsequently, the *mri_synthesize FreeSurfer* command was applied to create a synthetic FLASH image based on the previously calculated T1 (thresholded 1/R1 map) and proton density map. The optional flagged argument for optimal gray and white matter contrast weighting was used with the following parameters: 20, 30, and 2.5. Finally, the synthetic T1w image was divided by four according to the scale *FreeSurfer* expected. The pre-processing of the structural data using the *fMRIprep* toolbox was performed in the following steps: firstly, the synthetic T1w images were corrected for intensity non-uniformity (INU) with *N4BiasFieldCorrection* (Tustison et al., 2010), distributed with *ANTs 2.3.3* ((Avants et al., 2008), RRID:SCR 004757), and used as T1w-reference throughout the workflow. The T1w-reference was then skull-stripped with a *Nipype* implementation of the *antsBrainExtraction.sh* workflow (from *ANTs*), using OASIS30ANTs as target template. Brain tissue segmentation of cerebrospinal fluid (CSF), white-matter (WM) and gray-matter (GM) was performed on the brain-extracted T1w using *fast* (*FSL* 6.0.5.1:57b01774, RRID:SCR 002823, (Zhang et al., 2001)). Brain surfaces were reconstructed using *recon-all* (*FreeSurfer* 6.0.1, RRID:SCR 001847, (Dale et al., 1999)), and the brain mask estimated previously was refined with a custom variation of the method to reconcile ANTs-derived and FreeSurfer-derived segmentations of the cortical gray-matter of *Mindboggle* (RRID:SCR_002438, (Klein et al., 2017)). Volume-based spatial normalization to two standard spaces (MNI152NLin2009cAsym, MNI152NLin6Asym, where MNI stands for Montreal Neurological Institute) was performed through nonlinear registration with *antsRegistration* (*ANTs* 2.3.3), using brain-extracted versions of both T1w reference and the T1w template. The following templates were selected for spatial normalization: *ICBM 152 Nonlinear Asymmetrical template version 2009c* ((Fonov et al., 2009), RRID:SCR_008796; TemplateFlow ID: MNI152NLin2009cAsym), *FSL’s MNI ICBM 152 non-linear 6th Generation Asymmetric Average Brain Stereotaxic Registration Model* ((Evans et al., 2012), RRID:SCR_002823; TemplateFlow ID: MNI152NLin6Asym0.)

Voxel-Based Morphometry (VBM) data was derived from the synthetic T1w structural images via the standard SPM12 pipeline (https://www.fil.ion.ucl.ac. uk/spm/software/spm12/). This approach extracts spatially unbiased estimates of voxelwise GM volume. T1w images were automatically segmented into GM, WM, and cerebrospinal fluid and affine registered to the MNI template. A high-dimensional, nonlinear diffeomorphic registration algorithm (DARTEL) was used to generate a study-specific template from GM and WM tissue segments of all participants and then to normalize all segmented GM maps to MNI space with 2-mm isotropic resolution. All GM images were smoothed with a 4-mm full width at half maximum isotropic Gaussian kernel. Total brain volume was calculated by summing together the non-zero voxels in the modulated and warped GM and WM images of the VBM output (Malone et al., 2015).

### Functional MRI data pre-processing and connectome construction

First, a reference volume and its skull-stripped version were generated by aligning and averaging one single-band reference (SBRef). Head-motion parameters with respect to the BOLD reference (transformation matrices, and six corresponding rotation and translation parameters) were estimated before any spatiotemporal filtering using *mcflir*t (*FSL* 6.0.5.1:57b01774, (Jenkinson et al., 2002)). The estimated *fieldmap* was then aligned with rigid-registration to the target EPI (echo-planar imaging) reference run. The field coefficients were mapped on to the reference EPI using the transform. The BOLD reference was then co-registered to the T1w reference using *bbregister* (FreeSurfer) which implements boundary-based registration (Greve & Fischl, 2009). Co-registration was configured with six degrees of freedom. First, a reference volume and its skull-stripped version were generated using a custom methodology of *fMRIPrep*. Several confounding time-series were calculated based on the *preprocessed BOLD*: framewise displacement (FD), DVARS and three region-wise global signals. FD was computed following Power (absolute sum of relative motions (Power et al., 2014)). FD and DVARS are calculated for each functional run, both using their implementations in *Nipype* (following the definitions by Power et al., (2014)). The three global signals were extracted within the CSF, the WM, and the whole-brain masks. Additionally, a set of physiological regressors were extracted to allow for component-based noise correction (*CompCor* (Behzadi et al., 2007)). Principal components were estimated after high-pass filtering the *preprocessed BOLD* time-series (using a discrete cosine filter with 128s cut-off) for the two *CompCor* variants: temporal (tCompCor) and anatomical (aCompCor). For aCompCor, three probabilistic masks (CSF, WM and combined CSF+WM) are generated in anatomical space. The implementation differs from that of Behzadi et al. (2007) in that instead of eroding the masks by 2 pixels on BOLD space, the aCompCor masks are subtracted from a mask of pixels that likely contain a volume fraction of GM. This mask is obtained by dilating a GM mask extracted from the FreeSurfer’s *aseg* segmentation, and it ensures components are not extracted from voxels containing a minimal fraction of GM. Finally, these masks are resampled into BOLD space and binarized by thresholding at 0.99 (as in the original implementation). Components are also calculated separately within the WM and CSF masks. For each CompCor decomposition, the *k* components with the largest singular values are retained, such that the retained components’ time series are sufficient to explain 50 percent of variance across the nuisance mask (CSF, WM, combined, or temporal). The remaining components are dropped from consideration. The head-motion estimates calculated in the correction step were also placed within the corresponding confounds file. The confound time series derived from head motion estimates and global signals were expanded with the inclusion of temporal derivatives and quadratic terms for each (Satterthwaite et al., 2013). Frames that exceeded a threshold of 0.5 mm FD or 1.5 standardized DVARS were annotated as motion outliers. The BOLD time-series were resampled into standard space, generating a *preprocessed BOLD run in MNI152NLin2009cAsym space*. Many internal operations of *fMRIPrep* use *Nilearn* 0.8.1 ((Abraham et al., 2014), RRID:SCR_001362), mostly within the functional processing workflow. For more details of the pipeline, see the section corresponding to workflows in *<u>fMRIPrep</u>*<u>’s documentation</u>.

For the streamlined application of additional denoising components and data-cleaning strategies within a single framework, we utilized rs-Denoise (Kliemann et al., 2022) (please see https://github.com/adolphslab/rsDenoise), an open-source Python-based pipeline. This pipeline involved several steps: (1) z-score normalization of the signal at each voxel; (2) removal of linear and quadratic trends with polynomial regressors; (3) utilization of *fMRIPrep’s aCompCor* parameters, to regress out five components derived from whole-brain mean signals; (4) utilization of translational and rotational realignment parameters and their temporal derivatives as explanatory variables in motion regression; (5) temporal filtering was performed with a discrete cosine transform (DCT) filter with a cutoff frequency of 0.008 Hz. Lastly, the pre-processed runs were smoothed using a 4-mm full-width at half maximum (FWHM) Gaussian kernel and concatenated along the time domain. Individual fMRI recordings were then parceled into 416 cortical and subcortical brain regions using the Melbourne Subcortex Atlas (Tian et al., 2020) (Schaefer2018, 400 Parcels and 7 Networks and Tian Subcortex scale 1), and functional connectivity (FC) matrices were generated for each participant.

### Diffusion MRI data pre-processing and white-matter microstructure estimation

The preprocessing of dMRI data was executed using custom *MATLAB* scripts tailored in-house. These scripts proficiently filtered noise and eradicated prevalent artifacts such as Gibbs ringing, susceptibility distortion, motion, and eddy current-induced distortions. To provide further detail, data are denoised through the process of decomposition, which assumes that the variation occurring in the b-directions is similar in the neighborhood of the voxel. The method was adapted from Veraart et al. (2016). Gibbs ringing is corrected using the function ‘unring’, which is based on the approach described by Kellner (Kellner et al., 2015). *FSL*’s function *’eddy’* (http://fsl.fmrib.ox.ac.uk/fsl/fslwiki/EDDY) is an integrated approach correcting for off-resonance effects and subject movement in dMRI, and the methodology entails the following steps: first, *FSL*’s *’topup’* is employed to estimate the susceptibility field and generate unwarped b=0 images. Subsequently, the unwarped b=0 images are brain-masked using *FSL ‘bet’*. Finally, a combined eddy current correction, unwarping, and motion correction are performed using *FSL ‘eddy’.* Individual voxelwise fractional anisotropy (FA), mean diffusivity (MD), and radial diffusivity (L1) maps were computed using *dtifit* within the FSL software package (Smith et al., 2004). These four DTI features were selected based on their ability to capture different aspects of white matter microstructure. For example, FA is a scalar value indicating the degree of anisotropy in water diffusion within a voxel, thus distinguishing directional orientation from isotropy; MD, another scalar value, reflects the average magnitude of water diffusion within a voxel and provides insight into the overall diffusion rate and structural properties of the tissue. Unlike MD, which provides information independent of direction, the first eigenvalue (L1) indicates the magnitude of diffusion along the primary direction, correlating with myelin structure or myelination. FA image processing involved a tract-based spatial statistics pipeline with registration to the *FMRIB58_FA* standard space. This was followed by the skeletonization of the mean group white matter and the projection of individual data onto the skeleton. The resulting mean skeleton image was thresholded at FA 0.2, with other DTI metrics (MD, L1) projected onto the FA skeleton using the tbss_non_FA option. Prior to integration into the subsequent data fusion model, all DTI data were standardized to 1 mm isotropic resolution.

### Modalities fusion analysis

We employed LICA (Groves et al., 2011; Llera et al., 2019) to integrate inter-participant variability shared across five features: gray matter volume (VBM), white matter microstructure (FA, MD, L1), and functional connectivity (FC). LICA is a Bayesian multimodal extension of the ICA model that allows for simultaneous factorizations across multiple data modalities, connecting them at the participant level through a shared mixing matrix that represents each participant’s contribution (one scalar value per participant) to each independent component. This technique provides, for each independent component (IC), a vector indicating the contribution (weight) of each modality and a spatial map per modality showing the extent of spatial variation (Beckmann et al., 2005). Considering our sample size and the recommendation that the model order be less than 25% of the sample size (Groves et al., 2012), we report results from a 63-dimensional factorization. Given our primary interest in multimodal components, and the fact none of the unimodel components correlated with DRF (Supplementary Table 1), we excluded any components where a single modality contributed more than 50% of the total variance (Kohn et al., 2021; Van Oort et al., 2023). Additionally, seven components were driven by a single participant, therefore, these components were not included in the correlation analysis (Supplementary Figure 2). To demonstrate the robustness of the factorization choice, different model order (60 and 65-dimensional factorizations) decompositions were also performed (Supplementary Figure 3, 4 and 5). For visualization purposes, the spatial maps were thresholded at |Z| > 3.0.

### Statistical Analyses

Following the methodology of Llera et al., (2019), we conducted a permutation test to determine the significant Spearman partial correlations between the subject loadings on the independent components, derived from LICA, and our measure of DRF, controlling for age, sex, IQ, and total brain volume. Multiple comparisons were addressed using FDR correction (p<0.05), according to Benjamini and Hochberg (1995). The analyses were performed in R, and a fixed random seed was used to ensure the reproducibility of our results.

## Results

### Study population and general results

Participants reported an average DRF of 2.17 times per week (*SD*=2.05) and an average WAIS-IV score of 112.05 (*SD*=9.86) (Figure 1A). There was no evidence of age (*rho*=-0.035, *p*=0.57), sex (*rho*=0.116, *p*=0.062), or IQ-related (*rho*=0.060, *p*=0.367) differences in DRF. Nevertheless, in order to align with the methodology employed in previous studies, sex, age, and IQ were controlled for in the analyses.

**Figure 1:**
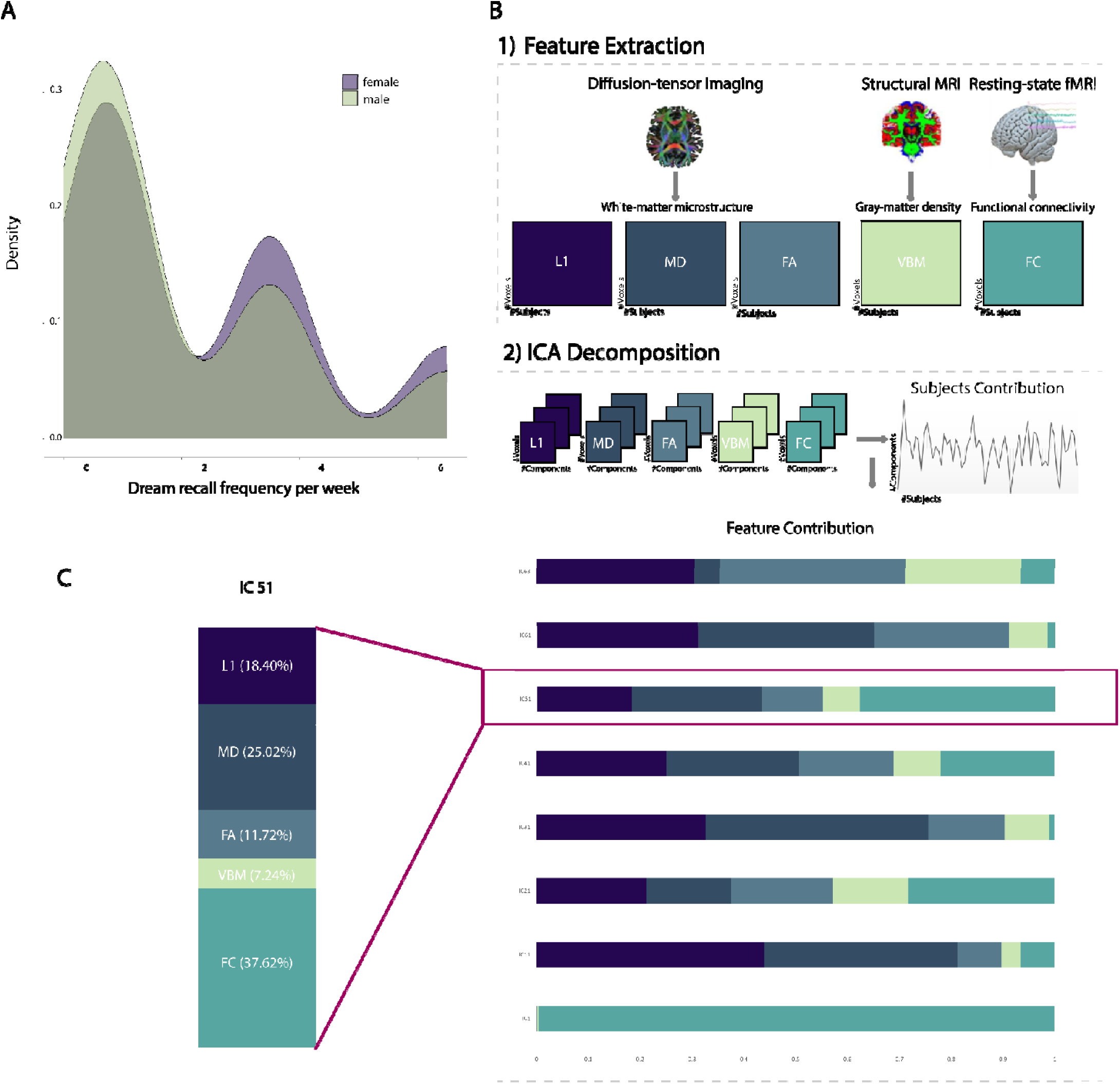
Demographic and LICA pipeline overview. A: Density distribution of the dream recall frequency scores (recoded into units per week) of females and males; B: (1) Diffusion-tensor, functional, and structural MRI data are used to extract relevant features, that is, radial diffusivity (L1), mean diffusivity (MD), fractional anisotropy (FA), gray matter volume as measured by Voxel-Based Morphometry (VBM), and functional connectivity (FC). (2) The aforementioned features are then utilized as input to the LICA algorithm, generating 63 independent components (IC), with the percentage of the distinct modalities contributions. Subsequently, the subject loadings of each independent component are combined with the behavioral data. (C) Among all independent components, multi-modal IC51 demonstrated a significant association with dream recall frequency.

### LICA decomposition and statistical results

LICA was used to decompose the multi-modal MRI data into 63 ICs (Figure 1B and Supplementary Figure 1). Of the 63 components, 46 were identified as multimodal, reflecting shared variance across different modalities. The statistical analysis revealed a single significant correlation between independent component 51 (IC51) and DRF (*rho*=-0.20, *p_FDRc_*=0.03), while controlling for total brain volume. To further confirm the stability of our results, we controlled for age, sex, and IQ in an additional partial correlation analysis (*rho*=-0.19, *p_FDRc_*=0.04). From the robustness analysis, we observed that IC51 is reproducible across different model orders (see, Supplementary Material for more details). The relative contributions from different modalities to IC51 were as follows: 18.40% for radial diffusivity (L1), 25.02% for mean diffusivity (MD), 11.72% for fractional anisotropy (FA), 7.24% for gray matter volume (VBM), and 37.62% for functional connectivity (FC) (Figure 1C).

Figure 2 presents the summarized images of each modality’s spatial map of IC51. DRF was associated with greater white microstructure integrity (reduced MD/L1 values) located in the frontal orbital cortex, parahippocampal gyrus, superior parietal lobule, and occipital cortex, particularly in the higher-order visual areas (V3 and V4). Furthermore, DRF was associated with lower gray matter volume in the occipital cortex (specifically in the V1 and V2 areas).

**Figure 2:**
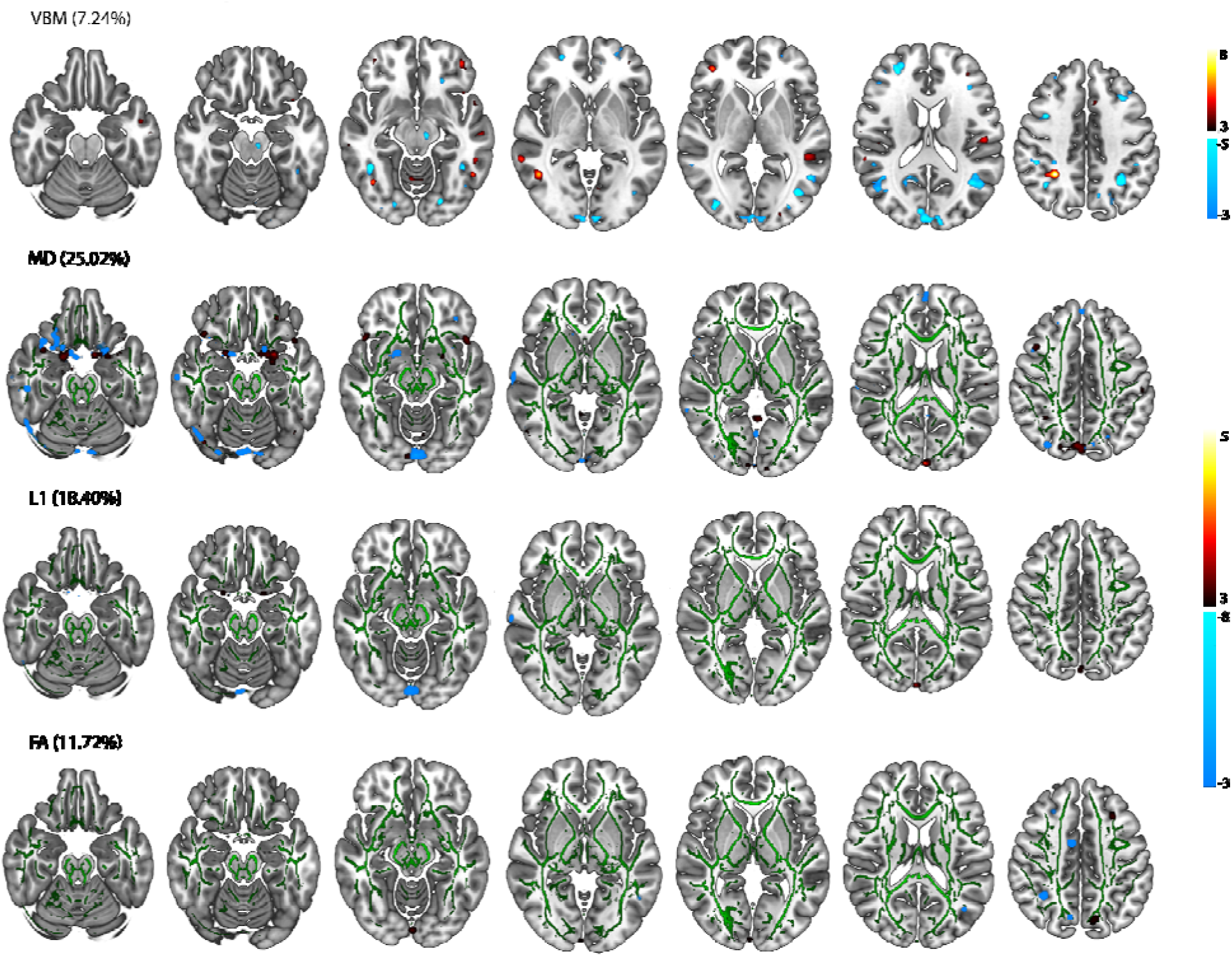
Brain Regions Associated with Dream Recall Frequency. Summary of the multimodal component (IC51) reveals the regions associated with dream recall frequency. The voxel-based morphometry (VBM) spatial map was thresholded at 3<|z|<8. The clusters of diffusion tensor imaging features were filled and thresholded at 3<|z|<8, then smoothed using a 0.3-mm Gaussian kernel in FSL for visualization purposes. Mean diffusivity (MD), radial diffusivity (L1), and fractional anisotropy (FA). The green map is the standard *FMRIB58_FA-skeleton* template provided in *FSL*.

Moreover, our results demonstrated enhanced functional connectivity within the occipital regions of the visual network, parietal regions of the default mode network, and sensorimotor networks (Figure 3A), and increased connectivity within the nucleus accumbens and left thalamus related to DRF. The observed relationships and the involvement of distinct brain regions underscore the complexity of the neural mechanisms underlying dream recall and emphasize the roles of microstructural and functional connectivity changes in this process.

**Figure 3:**
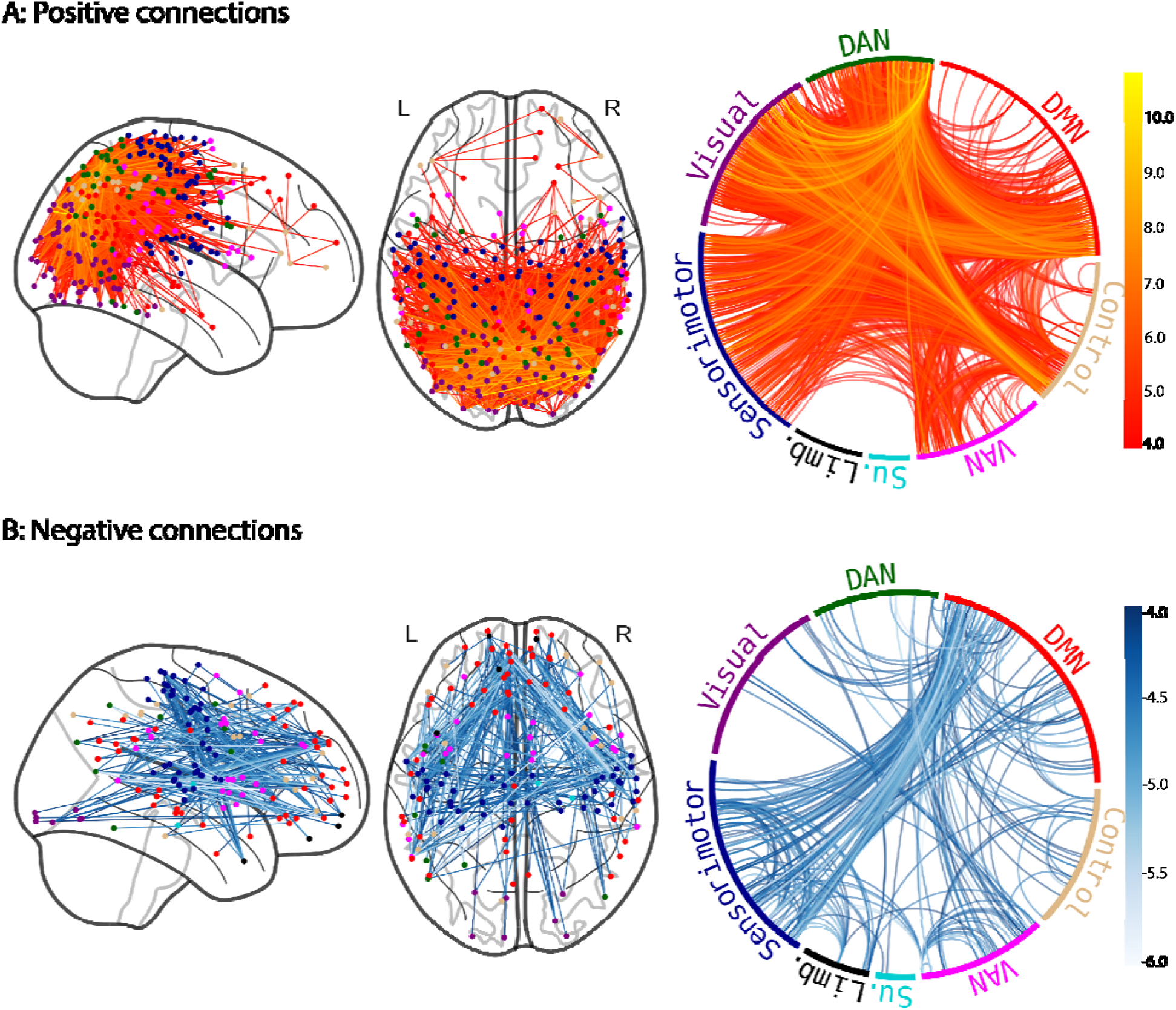
Inverse Relationship Between Functional Connectivity and Dream Recall Frequency. Functional connectivity is inversely associated with dream recall frequency. The connections were initially thresholded at |z|>3 and subsequently clustered according to their affiliation with the respective resting-state network. The positive (Figure 3A) and negative (Figure 3B) edges were thresholded at the 99th percentile for visualization purposes. DMN: Default mode network, Control: Control network, VAN: Ventral attention network, Su.: Subcortical network, Limb: Limbic network, Sensorimotor: Sensorimotor network, Visual: Visual network, and DAN: Dorsal attention network.

Additionally, the analysis demonstrated a reduction in functional connectivity, particularly between the parietal and temporal regions of the default mode, visual, sensorimotor, and dorsal attention networks, as well as within the dorsal attention network (Figure 3B). In contrast, DRF was associated with regions exhibiting increased FA and GM values, indicative of superior white matter microstructural integrity and gray matter volume. These regions included the middle frontal gyrus and several small clusters in the occipital and temporal cortex.

## Discussion

The present study employs a comprehensive, multimodal neuroimaging approach to investigate the neural correlates of DRF, with a particular focus on both brain structure and function. Our findings indicate an inverse relationship between DRF and brain microstructure integrity, volume, and functional connectivity. It is crucial to highlight that among the selected DTI modalities, high and low FA values indicate greater and poorer white matter microstructure, respectively. Conversely, for the MD and L1 modalities, high values indicate poorer microstructure integrity, whereas low values indicate greater white matter microstructure integrity. Our findings will be further interpreted in terms of their association between higher and lower DRF and the observed neuroimaging findings. For instance, we observed greater white matter microstructure integrity in several regions, including the frontal orbital cortex, parahippocampal gyrus, superior parietal lobule, and occipital cortex, particularly in the higher-order visual areas (V3 and V4) association with higher DRF. Furthermore, higher DRF was associated with lower gray matter volume in the occipital cortex (specifically in the V1 and V2 areas). Conversely, lower DRF was associated with reduced white-matter microstructure in the frontal orbital cortex, middle frontal gyrus, parahippocampal gyrus, and specific regions of the parietal cortex. These findings can be interpreted from a dual perspective: brain regions potentially contributing to dream generation and those related to DRF.

### Gray and white-matter morphology relationship with dream recall frequency

Dream experiences have been linked to localized increases in electroencephalogram (EEG) high-frequency (20–50 Hz) and reduced low-frequency (1–4 Hz) delta activity within posterior-occipital cortical regions during both REM and NREM sleep (Siclari et al., 2017). Similar patterns have been observed in dreams following NREM parasomnia episodes, where conscious experiences were associated with reduced delta and increased beta activity in the posterior cortical regions, including the primary visual cortices, occipital-temporal areas, medial temporal regions, and parts of the precuneus and posterior cingulate cortex (Cataldi et al., 2024). These findings suggest that dream generation is driven by distinct oscillatory patterns characterized by decreased low-frequency and increased high-frequency oscillations across specific brain areas, regardless of the sleep stage. Our findings align with these observations, as we observed enhanced white matter microstructure integrity in parietal-occipital regions, which are associated with higher DRF. This microstructure reflects well-organized and densely packed fibers that may facilitate optimal neural coordination and, thus, oscillatory activity. These results highlight the importance of particular brain areas and their microstructure integrity in facilitating the neural activity that underpins dream experiences and their frequency.

Our findings revealed a link between reduced gray matter volume in early visual areas (V1 and V2) and enhanced white matter microstructure integrity in higher-order visual areas (V3 and V4) and higher dream recall. In addition to processing fundamental visual characteristics such as color and pattern, V4 plays a role in visual learning, stimulus selection, and the translation of learned pattern relationships across the visual field. Furthermore, this area is modulated by attention, stimulus relevance, and perceptual context. V3 and V4 serve as critical connectors between early visual areas and higher-order cortical regions, integrating visual information across specialized channels and filtering it for higher-order brain regions (Farah, 1989). Empirical evidence supports the top-down model of dreaming, which proposes that cognitive processes, rather than sensory-motor inputs, primarily drive dream content (Foulkes & Domhoff, 2014). Studies have demonstrated that dreaming is associated with activity in higher-order brain regions, including the prefrontal cortex and association areas, which are crucial for imagination and narrative construction (Nir & Tononi, 2010). Increased high-frequency activity was observed in the frontal regions during NREM sleep when contrasting dream experiences with and without content recall (Siclari et al., 2017). This perspective is further reinforced by the observed greater microstructure integrity of the frontal orbital cortex, a region that integrates complex sensory information and is essential for processing reward values, learning associations, and emotional responses (Rolls, 2004). Our findings align with this perspective and support the idea that dreaming engages high-level cognitive processes, including those mediated by the frontal orbital cortex and higher-order visual areas. These results contribute to a more comprehensive model of dreaming, highlighting the importance of higher-order brain regions and cognitive systems in the formation and recall of dreams. The findings presented here can be extended to other brain regions, such as the parahippocampal gyrus, which plays a crucial role in connecting the default-mode network with the medial temporal lobe memory system and mediating functional connectivity between the hippocampus and posterior cingulate cortex (Ward et al., 2014). Furthermore, the parahippocampal gyrus plays a pivotal role in the relay of information between the hippocampal formation and other regions of the cerebral cortex, particularly the association cortices in monkeys (Van Hoesen, 1982), the direct electrical stimulation of the parahippocampal place area evoked topographic visual hallucinations, thereby demonstrating that the stimulation of higher-order visual areas can induce complex hallucinations in humans (Mégevand et al., 2014). Taken together, our findings indicate that individuals with greater white matter microstructure integrity in the parahippocampal gyrus recall their dreams more frequently, in line with these regions’ roles in processing contextual associations and memory processing. Given the methodological differences between our study and that of Zhou and colleagues (Zhou et al., 2019), a direct comparison is not possible. While Zhou et al. reported an inverse relationship between fiber integrity in the parahippocampal and fusiform gyri and DRF using a probabilistic tractography approach in an examination of 43 participants, we did not observe this same association in our data-driven analysis of white-matter integrity based on distinct DTI modalities. Their method specifically traced fibers connecting the two regions and focused on tracts that were consistently present across participants. In contrast, our approach did not explicitly assess fiber connectivity between regions. Furthermore, we did not identify any significant clusters in the fusiform gyrus, which makes direct comparisons with this previous finding challenging. Nevertheless, further research is required to more accurately define the spatial relationship between the parahippocampal area, dreaming, and trait dream recall.

Although we observed an association between higher DRF and greater white matter integrity in the frontal cortex, we did not find a link between increased medial prefrontal cortex white matter integrity linked to high DRF, as reported by Vallat and colleagues (2018). This discrepancy may be attributed to methodological differences between the studies. Our study employed DTI to assess white matter integrity, whereas Vallat et al. utilized voxel-based morphometry to quantify white matter density. Moreover, Vallat et al. focused on specific regions, including the medial prefrontal cortex, temporoparietal junction, hippocampus, and amygdala, comparing individuals with low and high DRF. In contrast, our approach was a whole-brain analysis that did not restrict the investigation to between-group comparisons. Instead, we examined regions associated with trait dream recall across a continuous spectrum. Additionally, although high DRF has been associated with increased rCBF in the temporoparietal junction (Eichenlaub et al., 2014), our observation of decreased white microstructural integrity in the inferior temporo-occipital gyrus, which has been associated with reduced rCBF (Chen et al., 2013), suggests a possible divergence from these results. These differences underscore the importance of methodological considerations and highlight the need for further research to reconcile these findings and fully understand the neural correlates of DRF.

### Functional connectivity relationship with dream recall frequency

Higher dream recall was associated with a widespread decrease in functional connectivity observed across various resting-state networks. This decrease was particularly evident between frontal, parietal and temporal regions of the default mode and visual networks, as well as between the sensorimotor and dorsal attention networks and within the dorsal attention network itself. A notable reduction in connectivity within the default mode network was observed, particularly in the right precuneus, prefrontal cortex, and posterior cingulate cortex. These findings are consistent with those of Zou and colleagues (2018), who reported a negative correlation between DRF and connectivity within the lateral visual network, the thalamus, and the posterior default mode network, thus indicating that decreased brain functional connectivity is linked to higher DRF. Similarly, we found that decreased functional connectivity in the thalamus, amygdala, globus pallidus, left hippocampus, and specific subregions of the visual, sensorimotor and dorsal attention networks was associated with frequent dream recall. In terms of functional connectivity relationship with high-frequency (20–50 Hz) neural oscillations, a study using laminar fMRI found a negative correlation between beta power and interregional layer connectivity, indicating that increased beta power reflects reduced laminar-specific connectivity in the visual cortex. In contrast, gamma band activity did not show a relationship with laminar connectivity, suggesting that while gamma activity is associated with the strength of the BOLD signal in middle and superficial layers, it does not correlate with changes in laminar fMRI connectivity within and between brain regions (Scheeringa et al., 2023). Clinically, pathological high-frequency oscillations (>80 Hz) have been linked to decreased cortical functional connectivity during seizure initiation and propagation (Ibrahim et al., 2013). Together, these findings offer valuable insights for interpreting our results, as they suggest that dream experiences accompanied by content recall are characterized by heightened high-frequency power in medial and lateral frontoparietal areas, potentially reflecting distinct neural dynamics underlying the recall of dream content (Siclari et al., 2017).

In contrast, lower DRF was associated with increased functional connectivity in the occipital areas of the visual network, parietal regions of the default mode and sensorimotor networks, as well as in the nucleus accumbens and left thalamus. Although our results are based on data collected during wakefulness, the increased functional connectivity within these regions may reflect underlying neural activity that supports low-frequency oscillations during sleep. Prior research has demonstrated a close relationship between low-frequency electrophysiological signals, such as delta oscillations, and resting-state fMRI signals. Specifically, the BOLD hemodynamic response has been shown to correlate with power coherence in the low-frequency delta band across various states of consciousness, including wakefulness, REM sleep, and NREM sleep, in both human and animal studies (Lu et al., 2007; He et al., 2008; Wilson III et al., 2016). The increased connectivity in posterior parietal, occipital, and thalamic regions observed in our study may indicate a stable neural configuration that is optimal for delta oscillation synchronicity and propagation. This hypothesis aligns with the neural dynamics observed during sleep, particularly when dream content recall is low, where increased delta oscillations are associated with diminished cortical activation and reduced conscious awareness and, consequently, dream experiences (Siclari et al., 2017). Although drowsiness and sleep-like activity can be observed during resting-state fMRI (Tagliazucchi & Laufs, 2014), the total recording time in our study was 18 minutes, shorter than typical resting-state or task-based recordings. While it is possible that some participants experienced brief periods of drowsiness, it is unlikely that most would have reached deeper sleep stages, where delta activity dominates. Instead, the association between resting-state functional signals and delta oscillations across various states of consciousness provides a plausible mechanistic explanation for our findings that increased functional connectivity observed in the parietal, occipital, and thalamic regions during wakefulness may serve as a precursor to the neural dynamics that occur during sleep, where local increases in delta power have been correlated with the absence of dream reports (Siclari et al., 2017). Further research is required to elucidate the relationship between functional connectivity and neural oscillations across different states of consciousness in humans.

### Limitations and conclusions

In the present study, we employed LICA to explore the neural correlates of trait dream recall. While LICA is an effective method for integrating data from different modalities, providing a comprehensive and biologically informative view of complex phenomena, several limitations should be noted. First, the efficacy of the method may be affected by variability in the number of features and distributions across modalities. Moreover, it is important to interpret the results of correlational studies cautiously, as there is currently no causal evidence to suggest that specific brain structure and functional features are directly involved in dream experiences. Our investigation of trait dream recall frequency may potentially overlook state components that may have influenced factors such as sleep stages, dream diaries, and daily events. These factors may have functional correlates rather than anatomical correlates. Future research may address these limitations by exploring additional measures related to state dream recall and sleep. The combination of simultaneous EEG/fMRI recordings over consecutive days, assessing both trait, retrospective and prospective dream recall, with serial awakening paradigms, has the potential to provide ongoing insights into the relationship between brain activity, dream production and dream recall. Furthermore, the incorporation of dream diaries would facilitate a more comprehensive capture of state-related aspects of DRF, given the potential for these to vary over time and influence the associations with anatomical and functional brain measures. Ultimately, validating our findings with neurostimulation techniques and extending the analysis to encompass both structural and functional brain aspects will be vital for a more comprehensive understanding of the neural correlates of dream memory recall. Further research is required to confirm these findings, with more diverse samples.

## Acknowledgments

We thank Dunja Paunovic, Katarina Vulic, Katarzyna Hat and Blanka Zana for their enormous efforts during data collection. This work was funded by the Dutch Research Council and the Bial Foundation.

## Notes

### Competing Interest Statement

The authors have declared no competing interest.

