## Supplementary Material for "The Neural Architecture of Dream Recall Frequency: Insights from Interindividual Variations in Brain Structure and Function"

* Shared senior authorship

*Corresponding author:*

Mariana Pereira

This supplementary material is intended to present the reader with further information about the analyses presented in the primary manuscript.

### Linked-Independent Component Analysis of Model Order 63

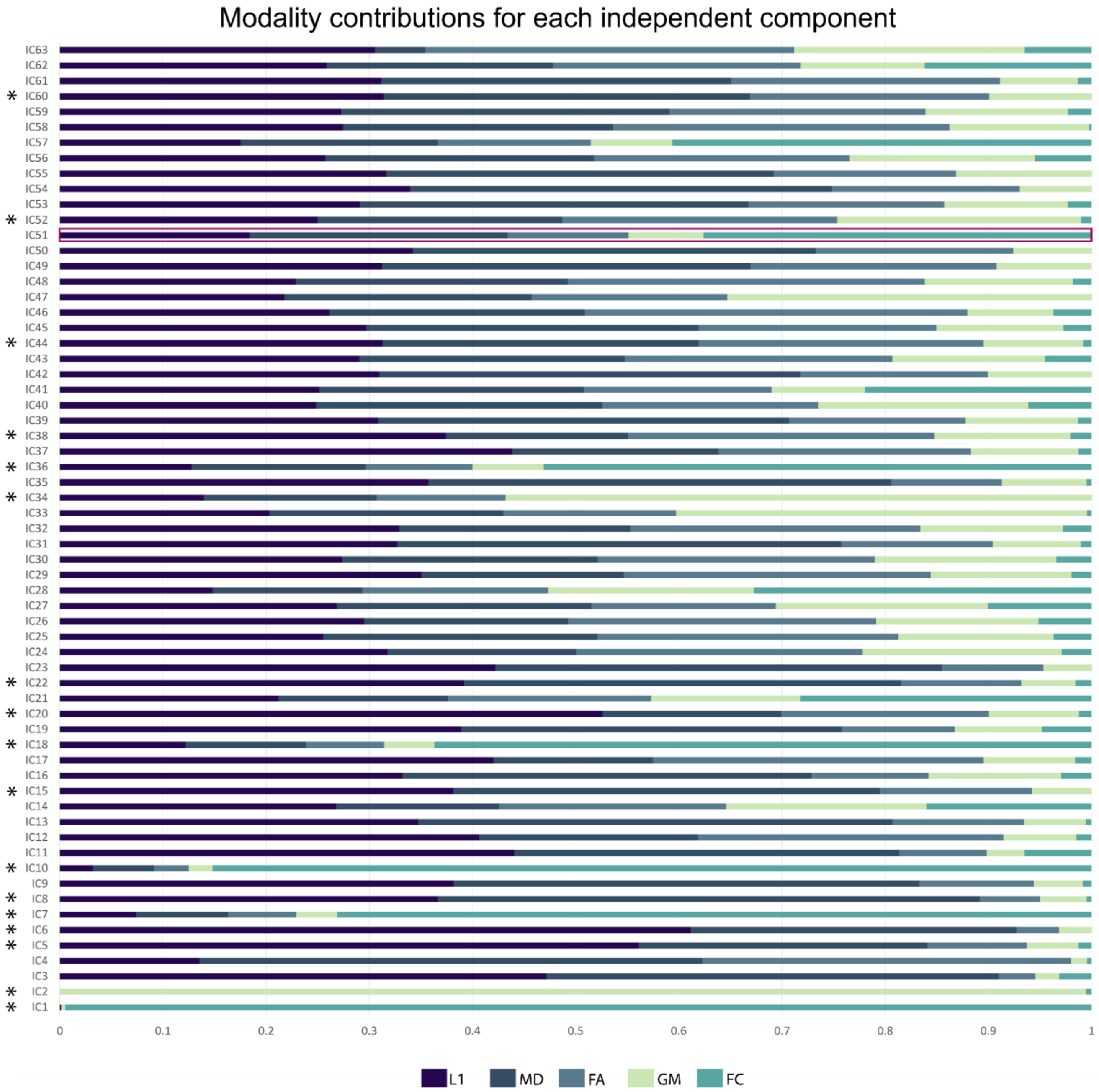


**Supplementary Figure 1:** Modality contributions for the 63-dimensional factorization. This study examines the modality contributions of the 63-dimensional factorization. The independent component highlighted in red significantly correlated with dream recall frequency. From the 63 independent components, those marked with an asterisk were excluded from the statistical analysis. This was because one single modality contributed to more than 50% of the total contributions, as was the case with components 1, 2, 5, 6, 7, 8, 10, 18, 20, 34, and 36.

Furthermore, components 10, 15, 22, 34, 38, 44, 52, and 60 were not included in the final statistical analysis. This was due to the fact that they were driven by a single subject. Radial diffusivity (L1), mean diffusivity (MD), fractional anisotropy (FA), gray matter volume (Voxel-Based Morphometry - VBM), and functional connectivity (FC).

**Supplementary Table 1:** Excluded unimodal independent components and their corresponding p-values uncorrected and corrected, respectively.

| **Excluded Unimodal Independent Components (ICs)** | | | |
|---|---|---|---|
| **IC** | **rho** | **p-value** | **p-value (corrected)** |
| IC1 | -0.0495 | 0.425 | 0.934642373 |
| IC2 | -0.09946 | 0.1177 | 0.934642373 |
| IC5 | 0.050721 | 0.4312 | 0.934642373 |
| IC6 | -0.02418 | 0.6971 | 0.934642373 |
| IC7 | -0.03533 | 0.5811 | 0.934642373 |
| IC8 | 0.091886 | 0.1408 | 0.934642373 |
| IC10 | -0.04008 | 0.5271 | 0.934642373 |
| IC18 | -0.05805 | 0.3484 | 0.934642373 |
| IC20 | 0.048375 | 0.4464 | 0.934642373 |
| IC34 | 0.043837 | 0.4974 | 0.934642373 |
| IC36 | 0.038101 | 0.5464 | 0.934642373 |

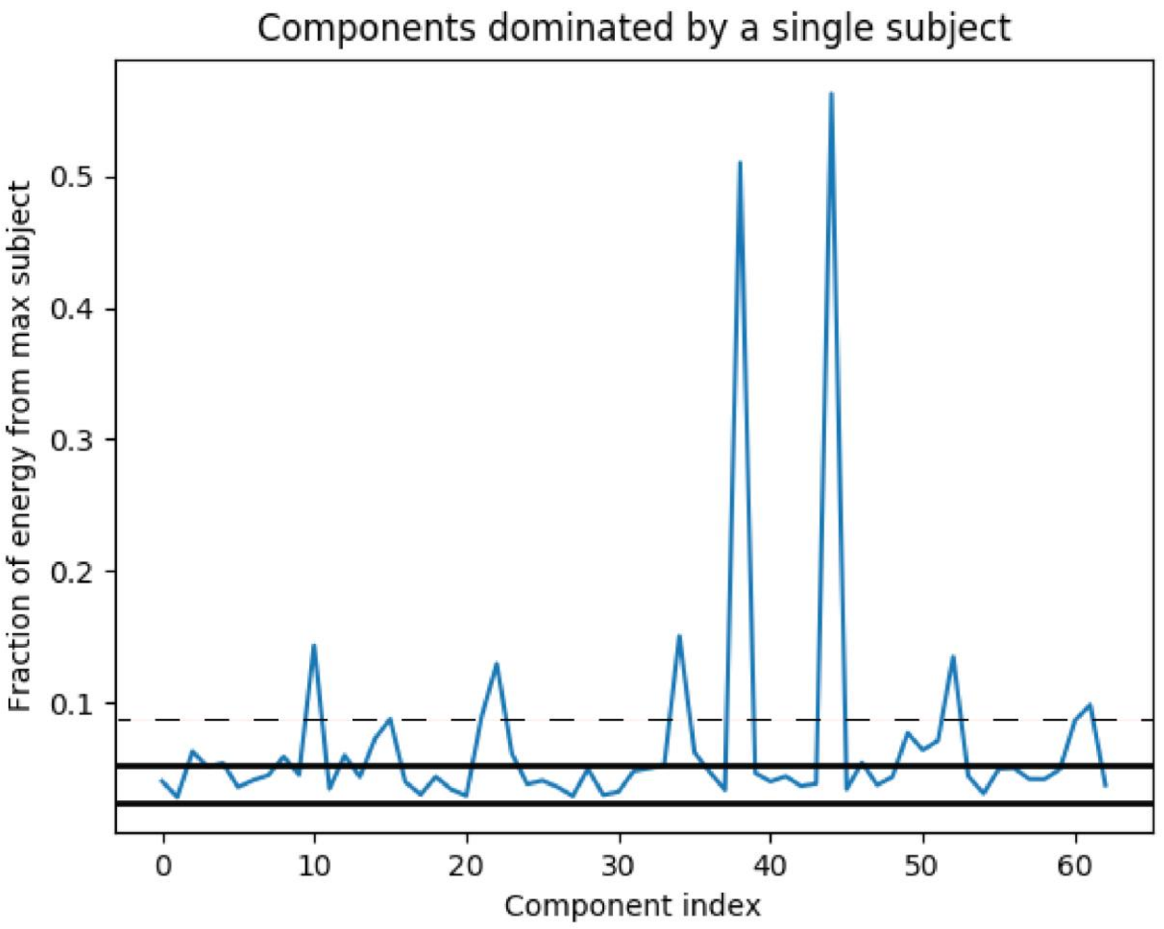


**Supplementary Figure 2:** The independent components dominated by a single subject, information output from LICA. Based on the fraction of energy, and represented by the dashed line, the following components were excluded from the statistical analysis: components 10, 15, 22, 34, 38, 44, 52, and 60.

**Robustness of the model order**

To evaluate the stability of the results obtained when different model orders are selected. Correlation analyses were conducted between the subject-mode components of the presented 63-dimensional factorization and those of the 60- and 65-dimensional factorizations. The top row of Figure 1 depicts the correlation matrices between the 63-dimensional factorization (yaxis) and the factorizations with 60 and 65 components (left and right panels, respectively). Only those correlations that were statistically significant after the false discovery rate (FDR) correction are displayed; that is, those with p-values smaller than 0.05/(63 × 60) and 0.05/(63 × 65). Moreover, the bottom row of Figure 1 illustrates the reproducibility of independent component 51. The figure depicts the sorted absolute correlations for IC51 across the model orders (60 vs. 63 and 63 vs. 65), thereby demonstrating its stability across dimensionality choices. As illustrated in the plot, IC51 exhibits consistent high correlation values across different model orders, thereby underscoring its robustness and emphasizing its significance in our analysis.

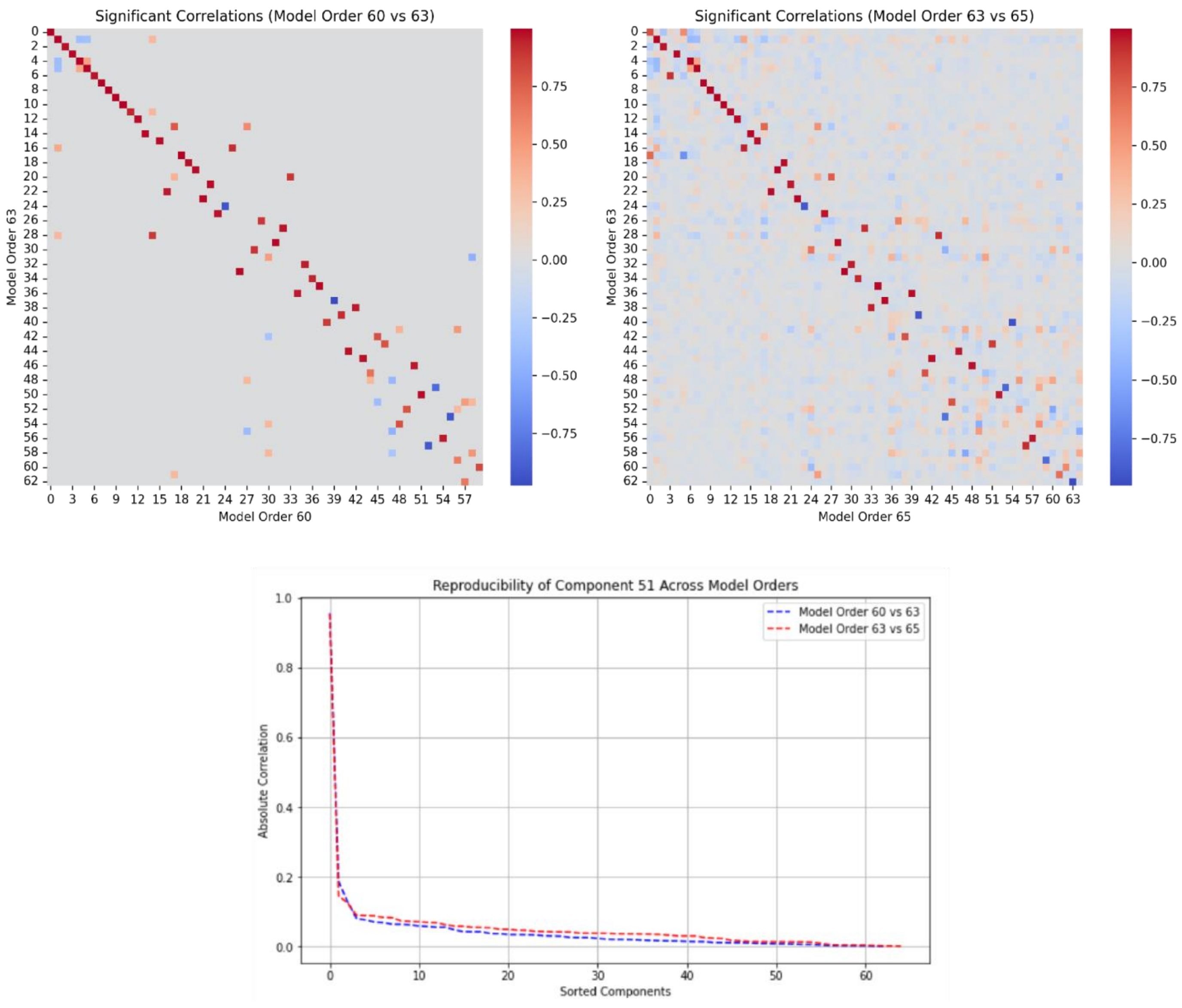

**Figure 3: Correlation Stability of Independent Component 51 Across Model Orders:** Significant correlations are demonstrated between the reported 63-dimensional factorization and the 60-dimensional (left panel) and 65-dimensional (right panel) factorizations. The bottom row presents sorted absolute correlations for independent component 51 for each of the 63-dimensional factorizations with the corresponding components from the other model orders, thereby highlighting its stability and robustness across model orders.

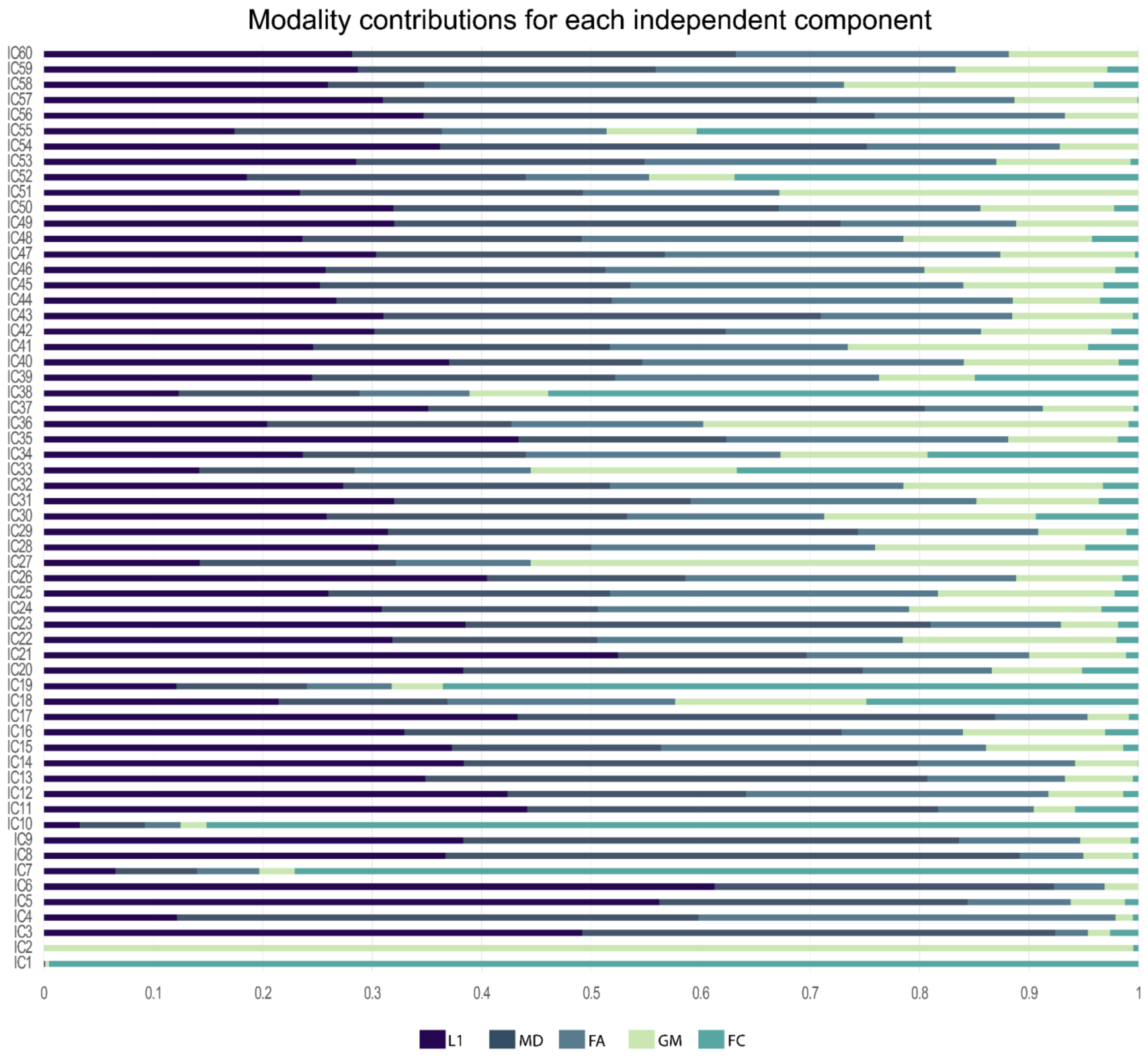


**Supplementary Figure 4:** Modality contributions for the 60-dimensional factorization.

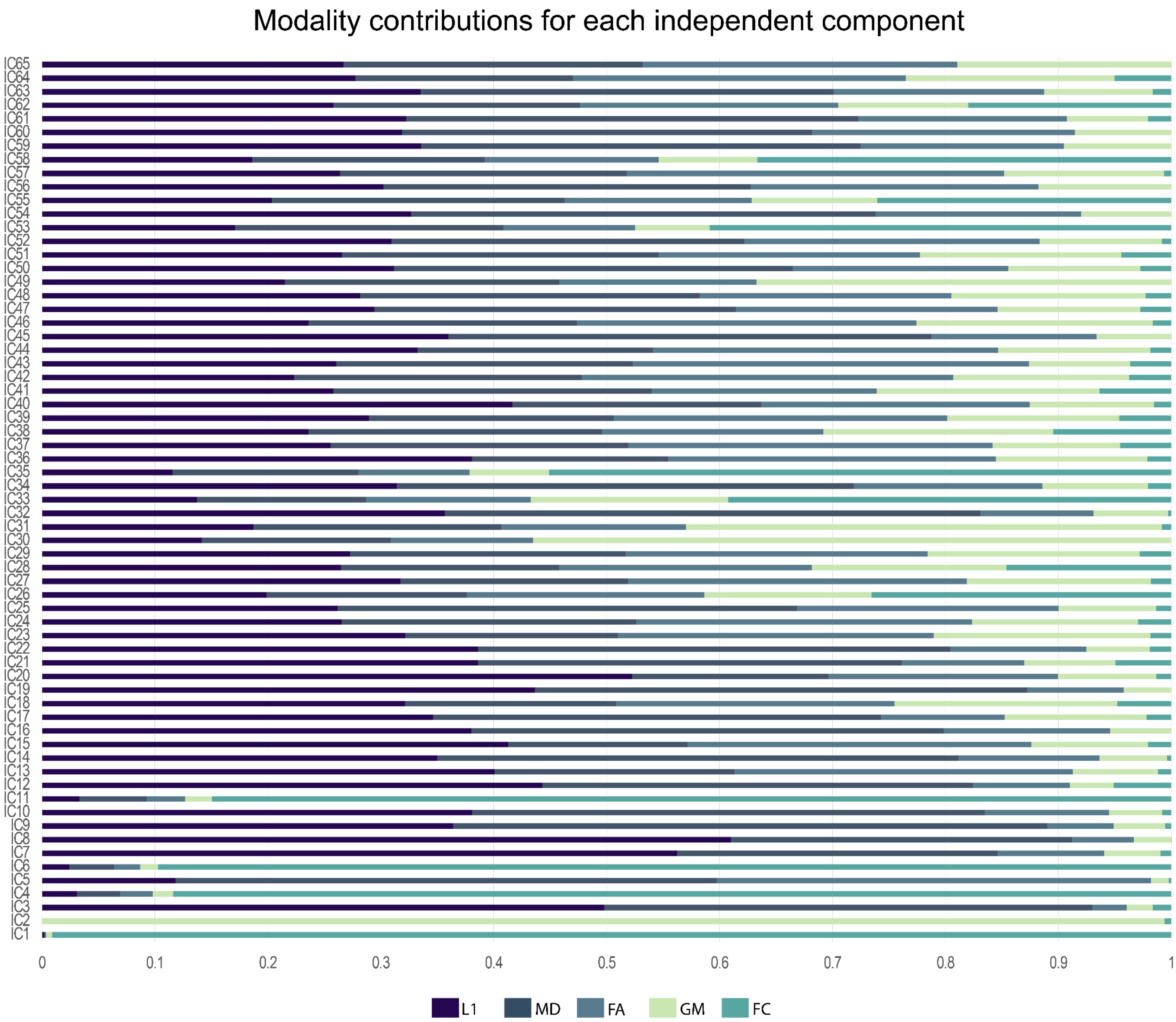


**Supplementary Figure 5:** Modality contributions for the 65-dimensional factorization.
